# Microsecond Molecular Dynamics Simulations and Markov State Models of Mutation-Induced Allosteric Mechanisms for the Light-Oxygen-Voltage 2 Protein : Revealing Structural Basis of Signal Transmission Induced by Photoactivation of the Light Protein State

**DOI:** 10.1101/2023.12.22.573121

**Authors:** Sian Xiao, Mayar Tarek Ibrahim, Gennady M. Verkhivker, Brian D. Zoltowski, Peng Tao

## Abstract

Avena Sativa phototropin 1 Light-oxygen-voltage 2 domain (AsLOV2) is the model protein of Per-Arnt-Sim (PAS) superfamily, characterized by conformational changes in response to external environmental stimuli. This conformational change is initiated by the unfolding of the N-terminal helix in the dark state followed by the unfolding of the C-terminal helix. The light state is defined by the unfolded termini and the subsequent modifications in hydrogen bond patterns. In this photoreceptor, β-sheets have been identified as crucial components for mediating allosteric signal transmission between the two termini. In this study, we combined microsecond all-atm molecular dynamics simulations and Markov state modeling of conformational states to quantify molecular basis of mutation-induced allostery in the AsLOV2 protein. Through a combination of computational investigations, we determine that the Hβ and Iβ strands are the most critical structural elements involved in the allosteric mechanism. To elucidate the role of these β-sheets, we introduced 13 distinct mutations (F490L, N492A, L493A, F494L, H495L, L496F, Q497A, R500A, F509L, Q513A, L514A, D515V, and T517V) and conducted comprehensive simulation analysis. The results highlighted the role of two hydrogen bond Asn482-Leu453 and Gln479-Val520 in the observed distinct behaviors of L493A, L496F, Q497A, and D515V mutants. The comprehensive atomistic-level analysis of the conformational landscapes revealed the critical functional role of β-sheet segments in the transmission of the allosteric signal upon the photoactivation of the light state.

## Introduction

The Per-Arnt-Sim (PAS) superfamily is widely present across the kingdoms of life, members of which are often found in pathways that regulate responses to environmental change, including circadian response pathway.^1,2^ As a subset of the PAS superfamily, light, oxygen, or voltage sensing (LOV) domains are small, sensor modules found in proteins, which respond to environmental signals including light, redox potential, and oxygen to regulate various biological processes. *Avena Sativa* phototropin 1 Light-oxygen-voltage 2 domain (*As*LOV2) is a model protein to study LOV function and allostery. It contains a flavin mononucleotide (FMN) cofactor located in the center of the PAS fold. A large α-helical region of the *As*LOV2 C-terminal to the PAS fold is termed the Jα helix (**Figure 1**).^3^ *As*LOV2 undergoes an allosteric activation from its resting dark state upon irradiation with blue light. A covalent bond is formed between Cys450 side chain in the PAS fold and C(4a) of the FMN, resulting in the formation of cysteinyl-flavin adduct, referred to as the light state.^4^ This light activation also leads to a large conformational change in the overall protein structure, including the unfolding of the Jα helix. The reversion from the light state to the dark state for LOV domains is spontaneous and starts when irradiation ceases. The reversion process includes the thiol bond scission and conformational change back into the dark state, and occurs within seconds to hours, depending on the types of LOV domain.^5^ With its characteristic conformational changes upon light activation, AsLOV2 has been extensively studied as a model of designing optogenetic switches.^6–8^

**Figure 1.**
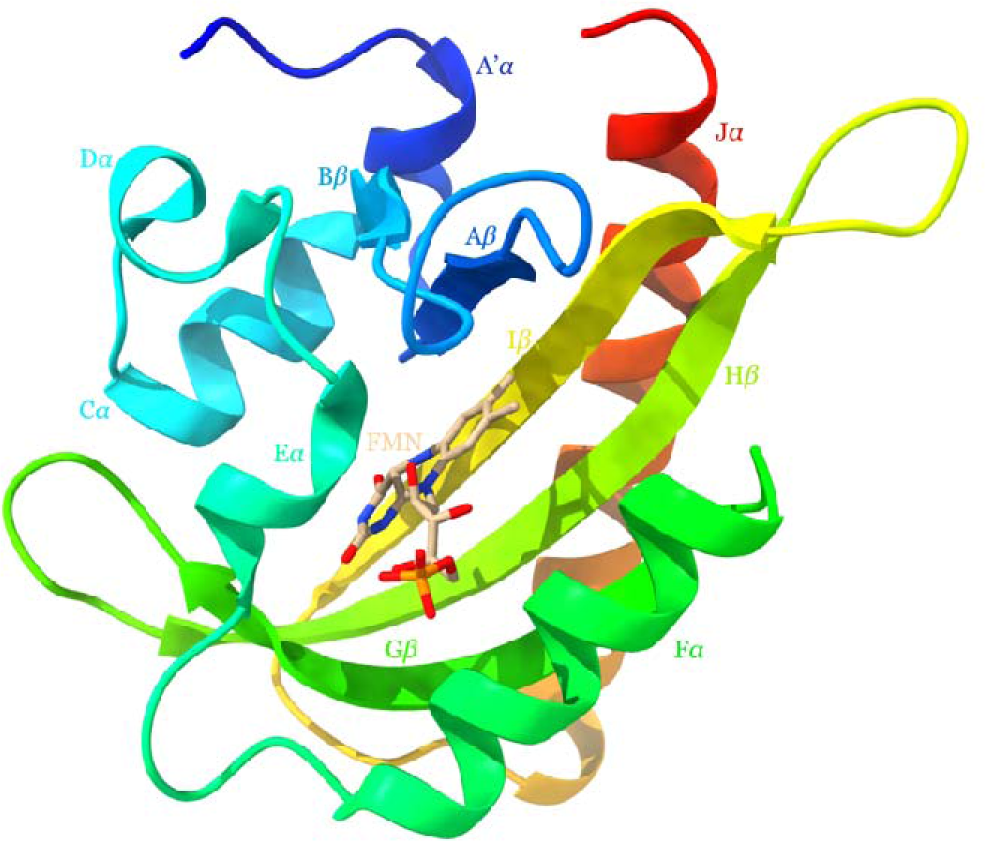
*As*LOV2 protein structure (PDB ID: 2V1B).

Past studies have identified the most characteristic conformational changes in *As*LOV2 to be localized in the termini, including the undocking and unfolding of the Jα helix at carboxyl terminal, and the unfolding of the A’α helix at amino terminal^9^. However, the whole signaling propagation pathway remains elusive with β-sheets being determined to play an important role in this process ^2,10,11^. Zayner *et al*. carried out an extensive mutational analysis of *As*LOV2 domain to study the relation between the sequence of the protein and its function, including 13 distinct mutations on Hβ sheet and Iβ sheet (F490L, N492A, L493A, F494L, H495L, L496F, Q497A, R500A, F509L, Q513A, L514A, D515V, and T517V)^12^. Photocycle time and conformational change of these mutations are analyzed but no obvious correlation exists between these two properties^12^. However, the ample experimental observations related to multiple mutations provide an opportunity for computational studies focusing on the allosteric mechanism underlying the activation process of the *As*LOV2 domain.

The crystal structures are mostly limited to proteins in their equilibrium states. Therefore, little information of the light activation of *As*LOV2 could be obtained directly from its crystal structures. Molecular dynamics (MD) simulations can capture the behavior of proteins in full atomic detail and at very fine temporal resolution and serve as important tools for understanding the physical basis of protein structure and function^13,14^. In this study, extensive MD simulations were conducted on the selected mutations of *As*LOV2 to investigate its allosteric mechanisms with the focus on the role of Hβ sheet and Iβ sheet. Combining simulation analysis and Markov state model^15,16^, we identified some intriguing mutant behaviors potentially due the change of the hydrogen bond pattern in mutants.

### Materials and Methods Molecular dynamics simulations

The initial structures of *As*LOV2 were obtained from the Protein Data Bank (dark structure: 2V1A, light structure: 2V1B)^17^. The covalent bond between C(4a) of FMN and Cys450 of *As*LOV2 was added in the light state. The 13 mutations listed in the Introduction section were introduced into the light states.

The simulation systems were prepared using CHARMM^18^ program package (version c41b1). Force field parameters for the FMN were obtained from a previous work^19^. The protonation states of titratable residues were determined under neutral pH. The protein systems were solvated in TIP3P^20^ water boxes with 10 Å buffer region on each dimension. In each system, sodium and chloride ions were added to neutralize the system and maintain an ionic strength around 0.15 M. The structures were minimized first with the steep descent method for 200 steps and the adopted basis Newton-Raphson minimization for 1000 steps afterwards. An initial 24 picosecond (ps) MD simulation was carried out to raise the temperature from 0K to 300K. The systems were subjected to 10 nanoseconds (ns) isothermal−isobaric (NPT) equilibrations at 300 K (equilibrium simulation), followed by 1 microsecond (μs) NVT-ensemble simulations (production simulation) at 300 K. Snapshots of product run were saved at 100 ps intervals. Three independent simulations were carried out for the wild type (WT) dark and light states of *As*LOV2 and the light states of 13 selected mutants. In all simulations, the SHAKE^21^ constraint was used to constrain the length of all bonds that involve a hydrogen atom. The water molecules are rigid during the simulation, with both their bond lengths and angles constrained. The long-range electrostatic interactions were accounted for using the particle mesh Ewald summation method^22^. The nonbonding interactions were cut off at 12 Å. The simulation was conducted using OpenMM^23^ (version 7.6.0).

### Simulation analysis

Root-mean-square fluctuation (RMSF) is a parameter to evaluate the flexibility of individual residues. *RMSF_i_* measures how much an individual residue *i* fluctuates around its average position 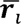 simulation with, frames. *RMSF_i_* is calculated as:

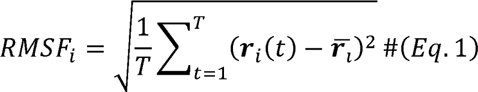

where *r_i_* (*t*) is the coordinate for residue, in frame L.

hydrogen atoms and acceptors (θ) and angles between donors, hydrogens, and acceptors (*r*)^24^. The donors Hydrogen bonds are identified by Baker-Hubbard method based on cutoffs for distances between

considered by this method are NH and OH, and the acceptors are O and N. The Hydrogen bonds analysis was carried out using MDTraj^26^ (version 1.9.7). The following criterion and the occurrence fractions during simulations were employed for validation of the hydrogen bonds:

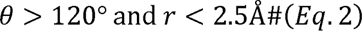

Dictionary of protein secondary structure (DSSP) is an algorithm used to assign standard secondary structure annotations to the amino acid residues of a protein.^25^ Based on the three-dimensional protein structure, it classifies residues as specific types of secondary structures, such as α-helices, β-sheets, turns, and bridges, based on the patterns of hydrogen bonding. The DSSP calculation is conducted by MDTraj using implementation of DSSP-2.2.0, giving the secondary structure assignments of each residue in each frame.

Dynamic cross correlation (DCC) analysis is a method used to analyze trajectories of MD simulations to detect correlative motions between residues or groups of residues within a molecule, which are important for understanding the function and interaction of the molecule. It involves calculating correlation coefficients between the movements of pairs of atoms over the course of a simulation, revealing how the motion of one part of a molecule is coupled to another. The DCC between the *q*th and *p*th atoms is defined by the following equation,

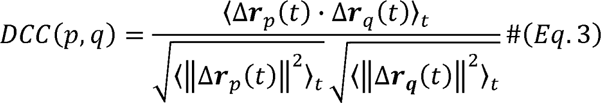

where *r_p_ (t)* denotes the vector of the *p*th atom’s coordinates as a function of time *t, …_t_* means the time ensemble average, and Δ **r***_p_* (*t*) = **r***_p_* (*t*)-, 〈**r***_p_*(*t*)〉*_t_* .

Essential dynamics of proteins propose that configurational space into two subspaces: an “essential” subspace containing only a few degrees of freedom, in which anharmonic motion occurs that comprises most of the positional fluctuations, and it describe motions which are relevant for the function of the protein; and the remaining space, named “physically constrained” subspace, in which the motion has a narrow Gaussian distribution and it merely describes local fluctuations.^27^ Essential dynamics analysis is performed using ProDy package^28,29^ (version 2.4.0).

### Markov State Model

The examination of MD simulations offers valuable insight into the conformational landscape and dynamic movements of biomolecules. However, the complexity and high dimensionality of the data from such simulations can obscure the crucial dynamic signatures^30^. To overcome this challenge and enhance the analysis of the conformational landscape, time-lagged Independent Component Analysis (t-ICA) is utilized for dimensionality reduction, projecting the extensive MD simulation data onto a more intuitive and interpretable low-dimensional space.

The t-ICA method identifies the slowest degrees of freedom and can preserve the kinetic information present in the MD trajectories. t-ICA finds coordinates of maximal autocorrelation at the given lag time. Therefore, t-ICA is useful to find the slow components in a dataset and thus an excellent choice as a dimensionality reduction method to transform MD data for the construction of a Markov state model.^31–33^

After obtaining the low-dimensional representation of the conformations from t-ICA analysis, k-means clustering method is conducted on projected low-dimensional space. Clusters identified in the clustering analysis is referred to as microstates. Each simulation frame is assigned to a microstate. Next step is to test whether these microstates are kinetically relevant and to look for an appropriate lag time τ. Relaxation timescales, also known as implied timescales, are examined against various lag time to generate a model satisfying the Markov assumption. After determining appropriate lag time, those microstates among which the simulation system could transit within the relaxation timescale are clustered together as one macrostate. The Perron-Cluster Cluster Analysis was used to identify macrostates, which are considered as kinetically separate equilibrium states^34,35^. More theory details can be found in a previous study^36^. The t-ICA and Markov state model building were conducted using the PyEMMA package^37^ (version 2.5.12).

## Results

### *As*LOV2 Wild Type

The root-mean-square fluctuation (RMSF) of each residue represented by alpha Carbon (Cα) was calculated for WT *As*LOV2 dark and light states, respectively. For each state, the reported RMSF values plotted in **Figure 2A** are averages across three independent simulations. RMSF based on each independent simulation of each state is provided in the Supporting Information (Error! Reference source not found.). Enhanced fluctuations are evident in the light state in both the A’α helix and Jα helix comparing to the dark state during the simulations, indicating an increase in flexibility within these regions upon light activation and the formation of the FMN-CYS450 covalent bond. Additionally, the RMSF is also elevated in the loop connecting the Hβ sheet to the Iβ sheet, the terminal residues of the Iβ sheet, and the loop between the Iβ sheet and Jα helix. The mean RMSF values among individual residues of A’α and Jα helices, Hβ and Iβ sheets were calculated respectively for further comparison (**Figure 2B**). The A’α helix at N-terminal shows the highest mean RMSF values among the selected secondary structures in both states. Furthermore, the A’α helix has a higher mean RMSF value in the light state than the dark state, indicating heightened mobility upon light exposure. The Jα helix follows this trend but to a smaller degree due to its larger size. The Hβ and Iβ sheets have lower mean RMSF values, with a slight elevation in the light state, denoting notable flexibility changes with light activation. This supports the conclusion that light activation induces a state of increased flexibility in specific regions of the protein, potentially facilitating its functional conformational changes.

**Figure 2.**
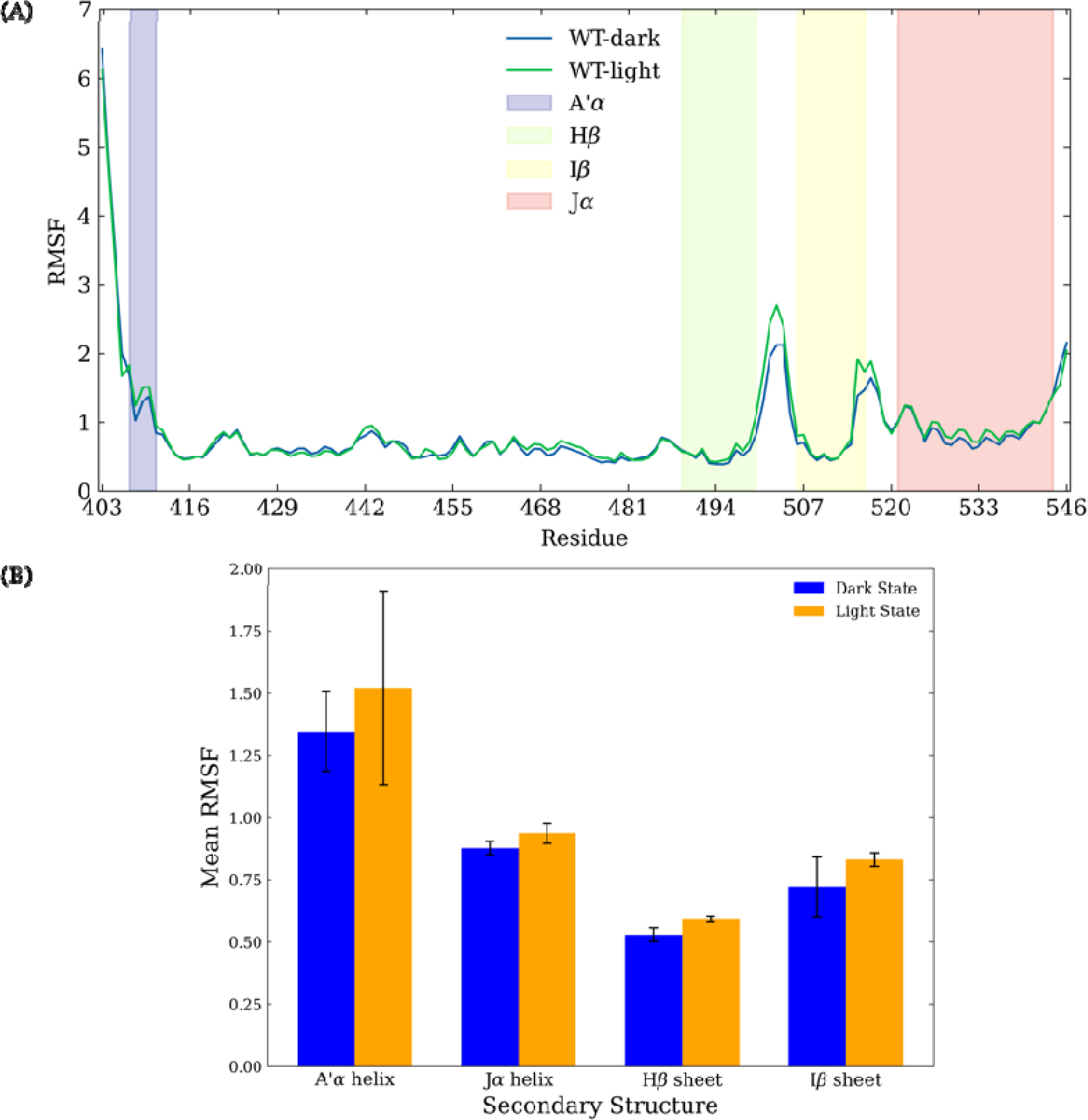
Root-mean-square fluctuations (RMSF) analysis of wild type (WT) of *As*LOV2 dark and light states. **(A)** RMSF of WT *As*LOV2 dark state (blue) and light state (green). The plotted RMSF is the averaged value over three independent simulations of each configuration. Four shadowed regions indicate the A’α helix (purple), Hβ sheet (green), Iβ sheet (yellow), and Jα helix (red), respectively. **(B)** Mean RMSF values of residues in four secondar structures for WT *As*LOV2 dark state (blue) and light state (orange), respectively.

Undocking and unfolding of both A’α and Jα helices are signature conformational changes during the transition from the dark state to the light state of *As*LOV2. Accordingly, the preservation of secondary structures in WT *As*LOV2 during simulations in both dark and light states is investigated. The ratio between the residues being recognized as correct structure and the total number of residues is calculated for each structure based on the simulation data of each state (**Figure 3**). For example, if a helix has ten residues, for a given simulation frame, eight residues in this helix are recognized with helical character using DSSP analysis, the preservation ration for this helix in this frame is 0.8. This is calculated and averaged for all frames. A significant decrease of the preservation ration of the A’α helix is evident suggests its unfolding during the simulation. Ratio analysis at the residue level for A’α helix is presented in Error! Reference source not found. and **Error! Reference source not found.**. In contrast, the Jα helix displays a stable helical ratio, with no significant changes between the two states. This suggests that the unfolding timescale of Jα helix exceeds the scope of conventional MD simulations. The Hβ and Iβ sheets exhibit a slight and moderate decrease in their sheet ratios, respectively. Given the typically stable nature of β-sheets, these observed decreases could be indicative of their involvement in the protein’s response to light activation, potentially playing a role in signal transduction. Combining with the RMSF analysis, these observations shed more light onto the protein conformation and dynamics change in response to light activation.

**Figure 3.**
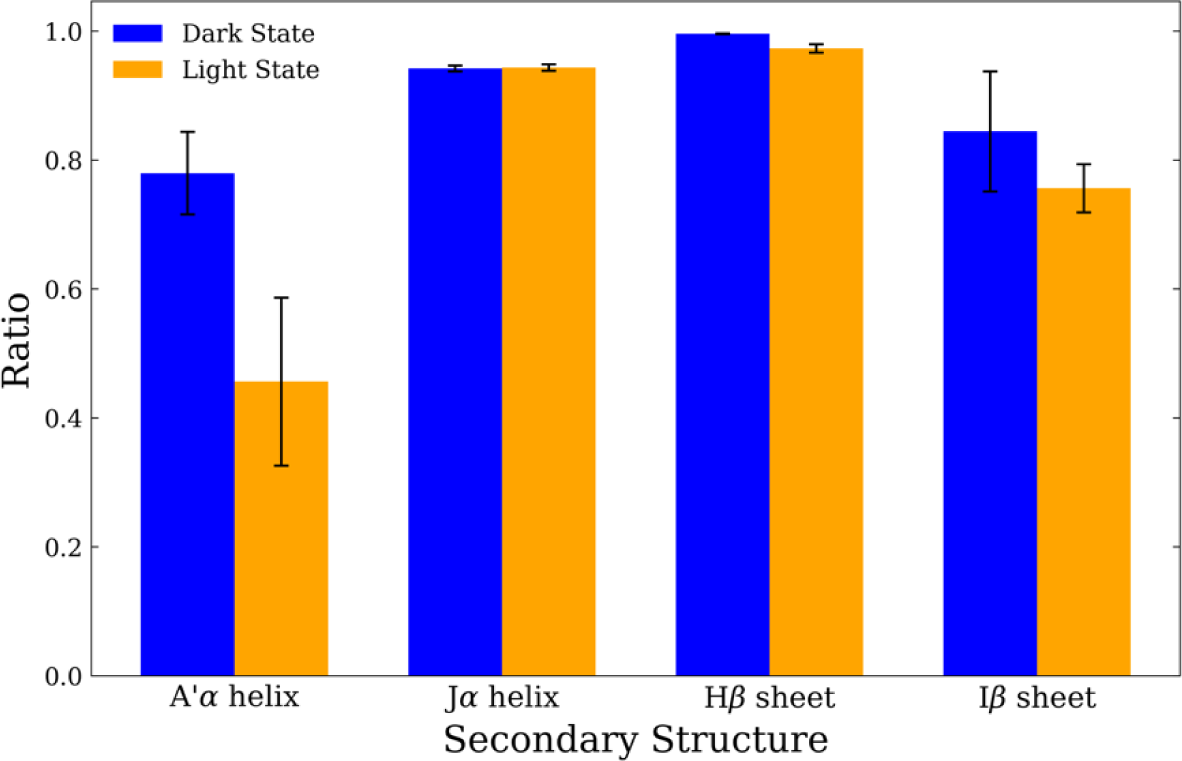
Average preservations of secondary structures for WT *As*LOV2 dark state (blue) and light state (orange) in four key secondary structures (A’α helix, Jα helix, Hβ sheet, and Iβ sheet).

The dynamical correlation within the dark and light states of WT *As*LOV2 is represented by the dynamic cross-correlation matrices (DCCMs) (**Figure 4**). For clarity, a discretized color scale with a segmentation threshold of 0.4 is used in the plots presented in **Figure 4** to accentuate the correlation contrasts between residues. This discrete segmentation enables the clear demarcation of highly correlated or anti-correlated regions. The DCCM plots with continuous color scale are provided in the Supporting Information (Error! Reference source not found.). In **Figure 4A**, the deep red hues along the diagonal show the robust positive correlation with neighboring residues, while the anti-diagonal also shows deep red, signaling a robust correlation among anti-parallel β-sheets. Upon light activation, as shown in **Figure 4B**, while the overall correlation structure remains similar, notable changes emerge in the pattern of Jα helix to the adjacent parts (specifically with anti-parallel β-sheet region and the Fα helix) of *As*LOV2 protein with increasing negative correlation. Such negative correlation is indicative of a potential undocking tendency, where the Jα helix exhibits a distinct movement away from the core structure in response to light activation. This tendency supports the light-induced undocking behavior of J_α_ helix, even though the undocking of the J_α_ helix was not observed in the current study.

**Figure 4.**
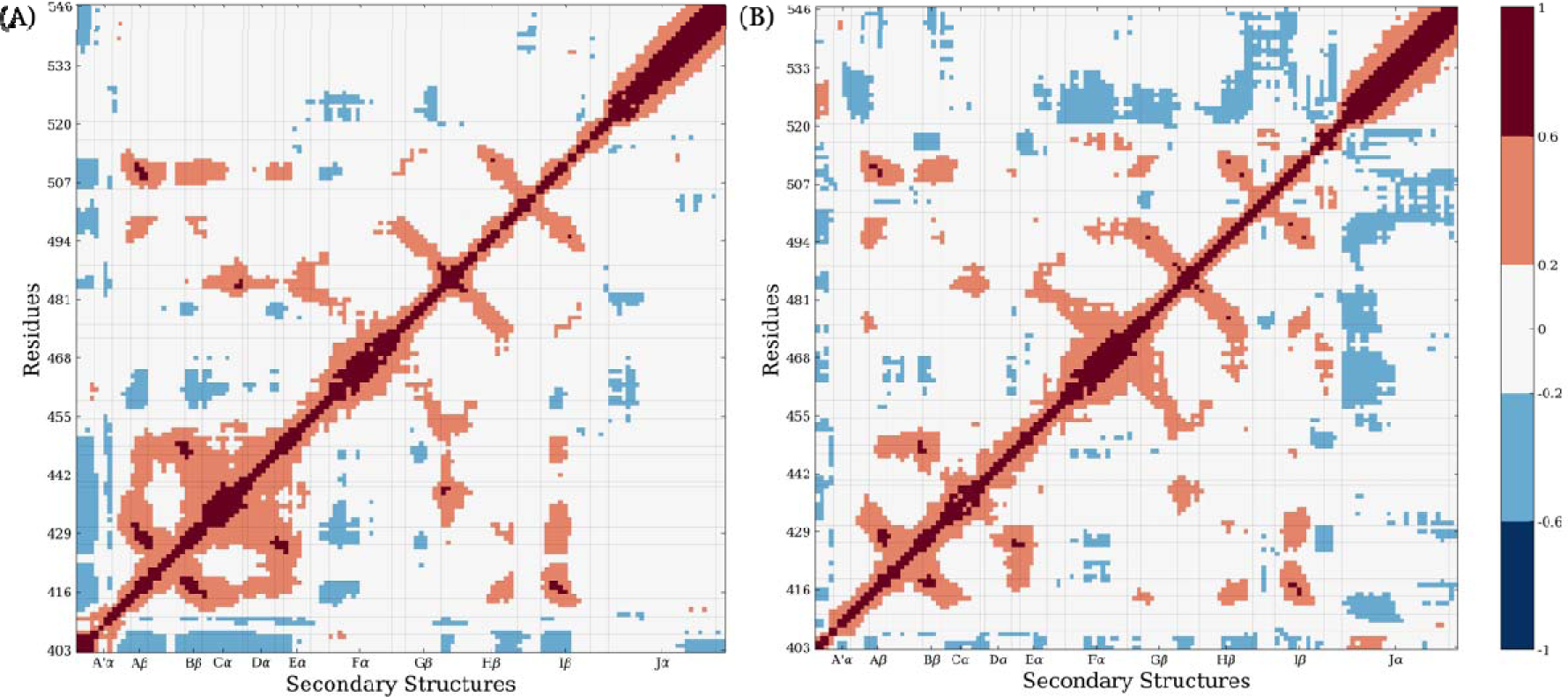
Discrete average dynamic cross-correlation matrix for WT *As*LOV2 in **(A)** dark state and **(B)** light state over three parallel simulations. For clarity, a discretized color scale with a segmentation threshold of 0.4 is used.

### *As*LOV2 Light States Mutants

Simulations of the light states of the selected *As*LOV2 mutants were carried out to compare the impact of the various mutants on the dynamics of *As*LOV2 domain. The comparative analysis of the light states of WT and mutants of *As*LOV2 focuses on the four key secondary structures: A’α and Jα helices, Hβ and Iβ sheets. The average RMSF values over residues of these four secondary structures in all light states are plotted in **Figure 5**. The RMSF plot for each mutant is shown in **Error! Reference source not found.**. The comparison of average RMSF values over the whole *As*LOV2 domain for each light state is shown in **Error! Reference source not found.**. The RMSF values of A’α helix vary significantly among all light states under investigation (**Figure 5A**). Large error bars suggest significant variability among the simulations. This could also be attributed to the short length of the A’α helix and its inherent flexibility given its location at the flexible N-terminal region. In contrast, the Hβ sheet, Iβ sheet, and Jα helix of most mutants exhibit RMSF values close to the WT value, indicating minimal impact from the corresponding mutants on protein flexibility. L493A, however, emerges as an outlier, exhibiting elevated RMSF values not just in these secondary structures but also throughout the entire protein (**Error! Reference source not found.**), implying a substantial increase in flexibility caused by the mutation. Additionally, the D515V mutation, located on the Iβ sheet, uniquely reduces the mean RMSF of the Iβ sheet itself, which suggests localized structural stabilization. These specific changes in dynamics necessitate additional investigation to elucidate the mutants’ impact on the protein’s behavior and interactions.

**Figure 5.**
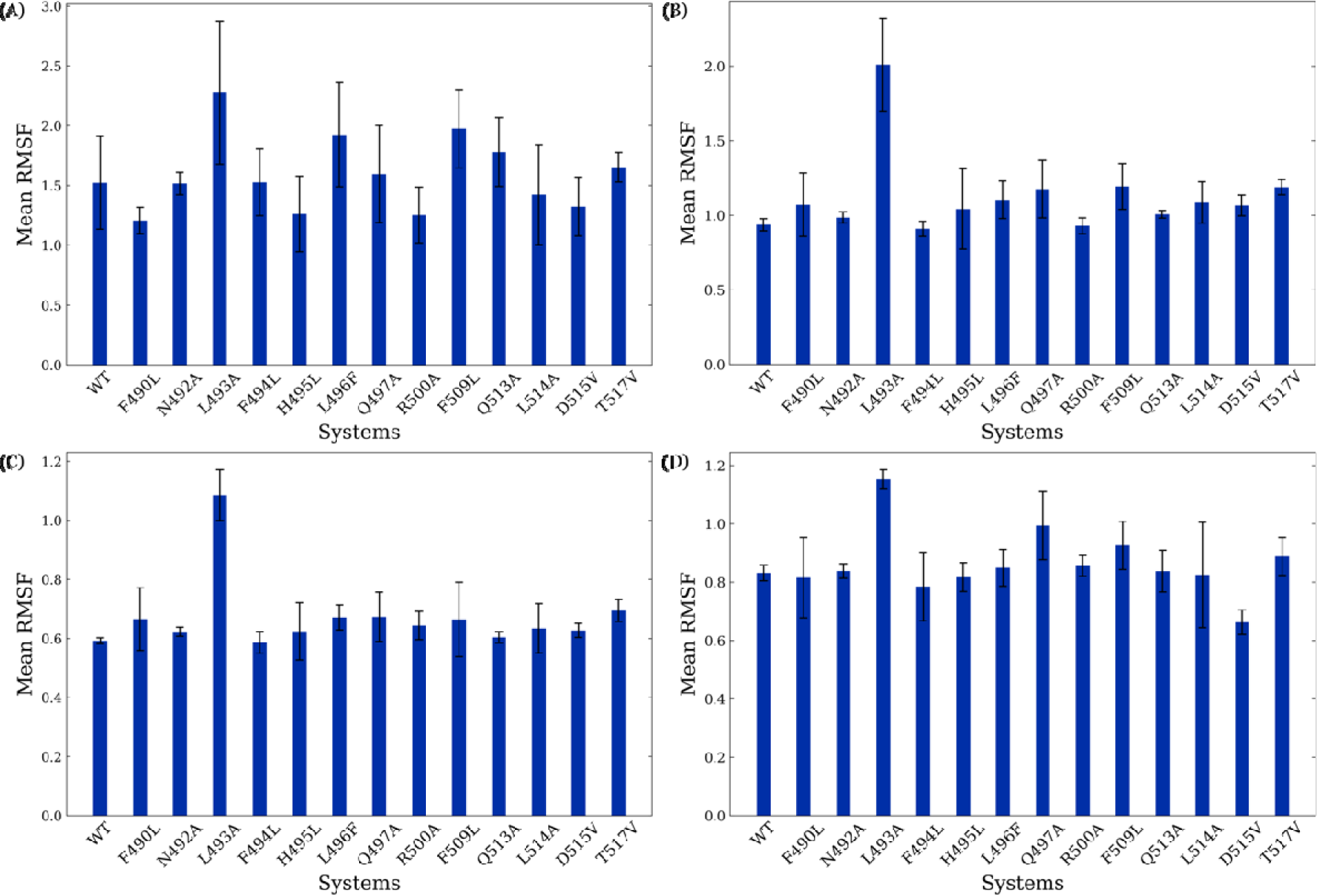
Average RMSF values over residues of WT and mutants in light state for **(A)** A’α helix, **(B)** Jα helix, **(C)** Hβ sheet, and **(D)** Iβ sheet.

The secondary structure preservations of the above four secondary structures in light states of *As*LOV2 WT and mutants are calculated and plotted for comparison (**Figure 6**). Consistency is observed in the Hβ sheet and Jα helix across all mutants, with the proportion of these structures close to that of the WT. The A’α helix, on the other hand, exhibits increased helical content in the light state among most mutants compared to the WT, yet still lower than the WT dark state values, suggesting a light-dependent partial unfolding or destabilization. While the A’α helix ratios show a general increase, the elongated error bars signal considerable variability, complicating definitive conclusions about the mutants’ influence. A general but not precisely negative correlation between helix ratio and RMSF underscores a correlated relationship between flexibility and unfolding behavior. Unlike a previous study focusing on mutants located on A’α helix itself, it is difficult to ascertain definitive conclusions about Hβ sheet and Iβ sheet mutants’ influence on the unfolding behavior of A’α helix. The Iβ sheet displays interesting responses to Q497A and D515V. Compared to other mutants, Q497A exhibits a higher Iβ sheet RMSF (**Figure 5D**) and a correspondingly lower Iβ sheet ratio (**Figure 6D**), suggesting an increase in flexibility. Conversely, D515V shows a lower Iβ sheet RMSF and a higher Iβ sheet ratio, implying enhanced stability. Therefore, Q497A and D515V are present as prime candidates for further investigation to deepen our understanding of their impact on the Iβ sheet’s dynamics.

**Figure 6.**
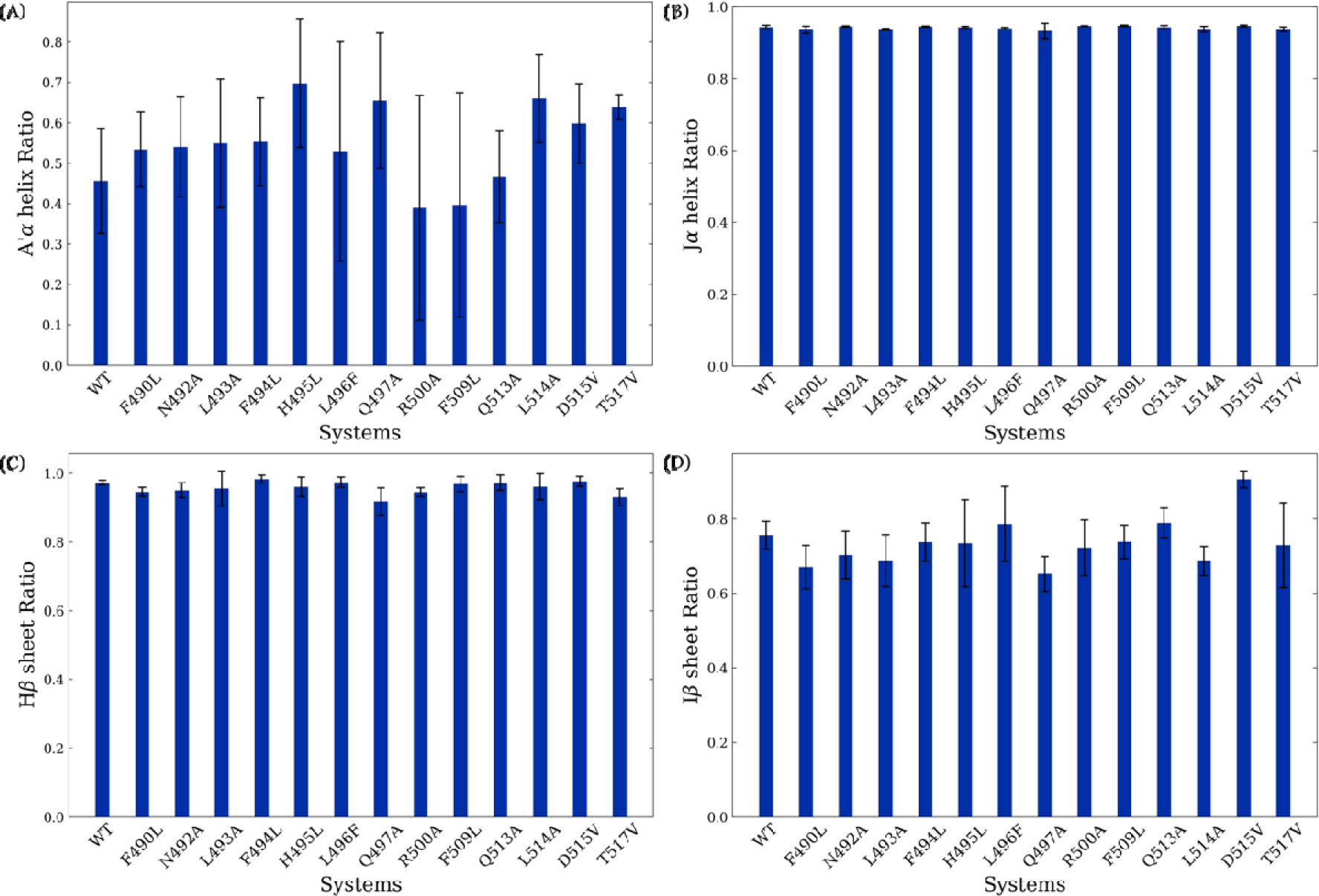
Average preservations of secondary structures for WT and mutants in light state for **(A)** A’α helix, **(B)** Jα helix, **(C)** Hβ sheet, and **(D)** Iβ sheet.

Dynamic cross-correlation analysis across various mutants consistently shows that the Jα helix maintains a negative correlation with the adjacent anti-parallel β-sheet region of the AsLOV2 protein. A subset of mutants demonstrates DCCM patterns remarkably akin to the WT with mutant R500A as an example (**Figure 7A**). Intriguingly, in several mutants including L496F (**Figure 7B**), a novel trend emerges wherein the Fα helix exhibits increasing negative correlations with spatially distant secondary structures: Aβ and Bβ sheets, as well as the Cα and Dα helices, despite being separated by the Eα helix and FMN. These mutants exhibit an uptick in negative correlations between inter-secondary structures, such as the loops connecting Fα helix and Gβ sheet and the loop connecting Gβ sheet and Hβ sheet. In F509L mutant, the C-terminal displays stronger negative correlations with the distant cluster comprising the Aβ and Bβ sheets as well as the Cα and Dα helices (**Figure 7C**).

**Figure 7.**
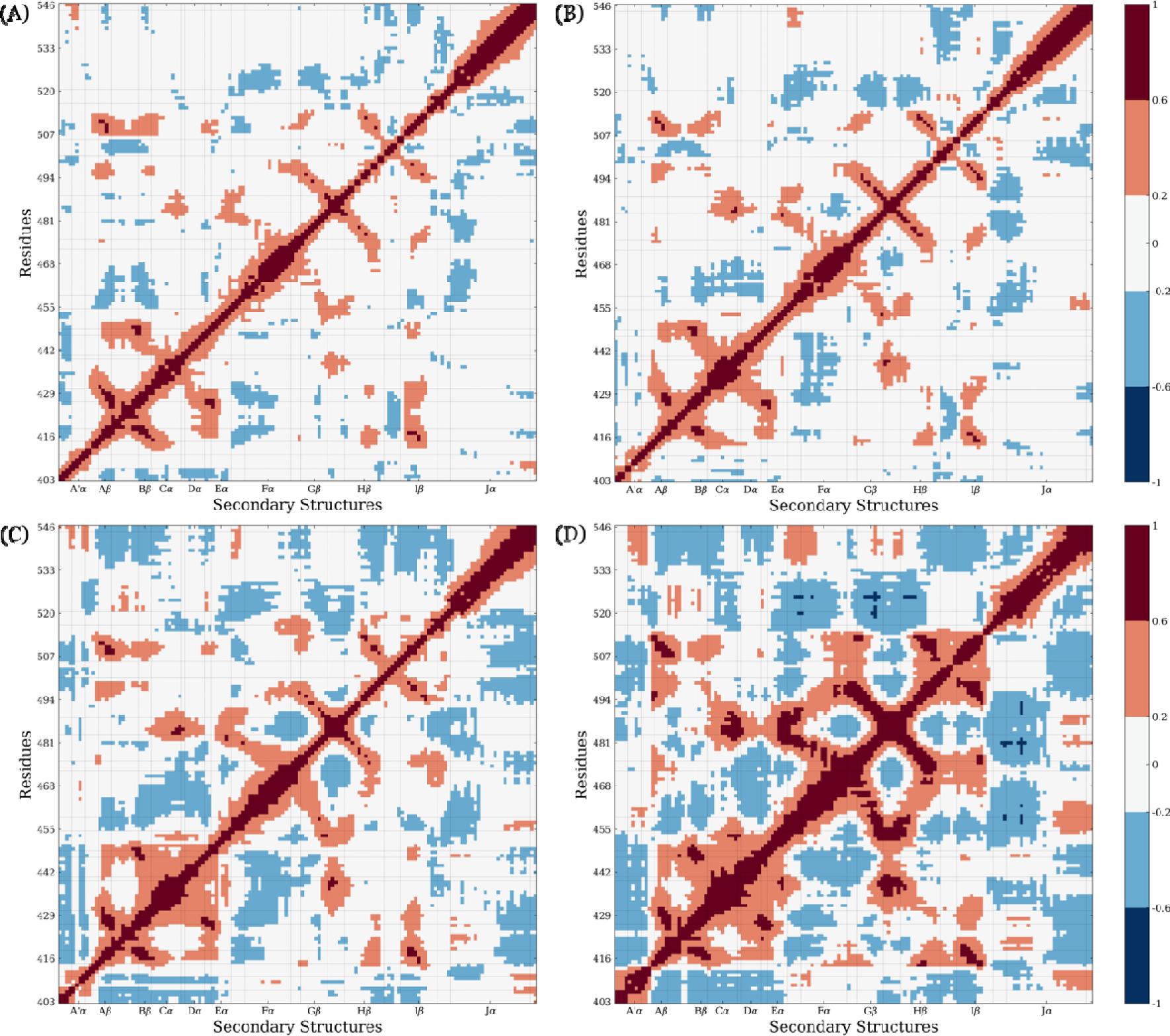
Representative discrete average dynamic cross-correlation matrix for light state **(A)** R500A, **(B)** L496F, **(C)** F509L, and **(D)** L493A.

An extreme case is presented by L493A, in which the cluster of Aβ and Bβ sheets, and the Cα and Dα helices, shows pronounced positive intra-group correlations. The anti-parallel β-sheets also display strong positive correlations with other β-sheets. Meanwhile, the Jα helix and the C-terminal reveal more negative correlations with these groups yet exhibit increased positive correlations with the A’α helix and Fα helix located in the same periphery (**Figure 7D**).

For each system in light state, including WT and mutants, top 10 essential modes are extracted and mean square-fluctuations of each residue in these 10 modes are calculated and plotted (**Figure 8**). The most significant contributions to these modes come from the following regions: N-terminal with A’α helix; the loop between Hβ sheet and Iβ sheet; the loop between Iβ sheet and Jα helix; and C-terminal with Jα helix. Most mutants exhibit higher flexibility compared to WT, especially at the loop between Hβ sheet and Iβ sheet, consistent with the RMSF results. L493A, H495L, and F509L show some extra essential movements at Cα helix (residues 431 to 439, spatially close to A’α helix) and the loop between Gβ sheet and Hβ sheet (residues 483 to 489, spatially close to Cα helix). Several mutants, including F490L, L493A, L496F, Q497A, F509L show higher flexibility at the loop between Iβ sheet and Jα helix. Especially, L493A extends the movement flexibility from this loop to the Jα helix.

**Figure 8.**
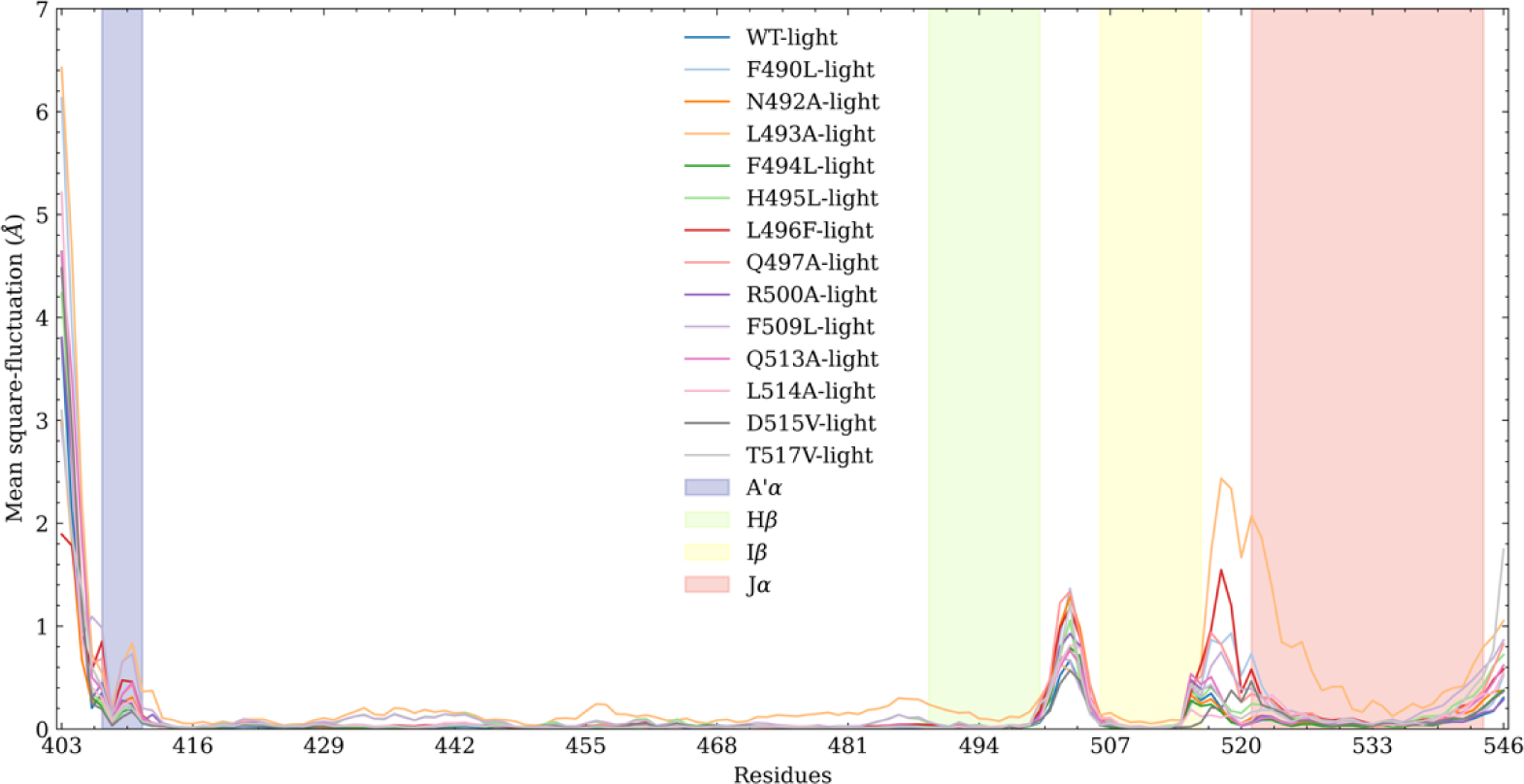
Mean square-fluctuations for top 10 essential modes for each system in light state.

### Dimensionality Reduction Analysis and Markov State Model

The simulations of all WT and mutant light states were combined and subjected to t-ICA. The simulations of each state are projected onto a two-dimensional (2D) space defined by the two vectors: IC 1 and IC 2. The mutants display drastically different distribution with L493A and L514 as the most significant cases (**Figure 9A**). The overall population of the combined simulations data is illustrated in **Figure 9B**. The WT distribution (not visible in **Figure 9A**) represents the most probable region represented as light yellow region in **Figure 9B**. Most mutant simulations resemble the WT distribution and exhibit slight deviation. Several mutants, including L493A, H495L, L496F, F509L, and L514A, exhibit a more pronounced deviation from the WT distribution and explore a much larger conformational space.

**Figure 9.**
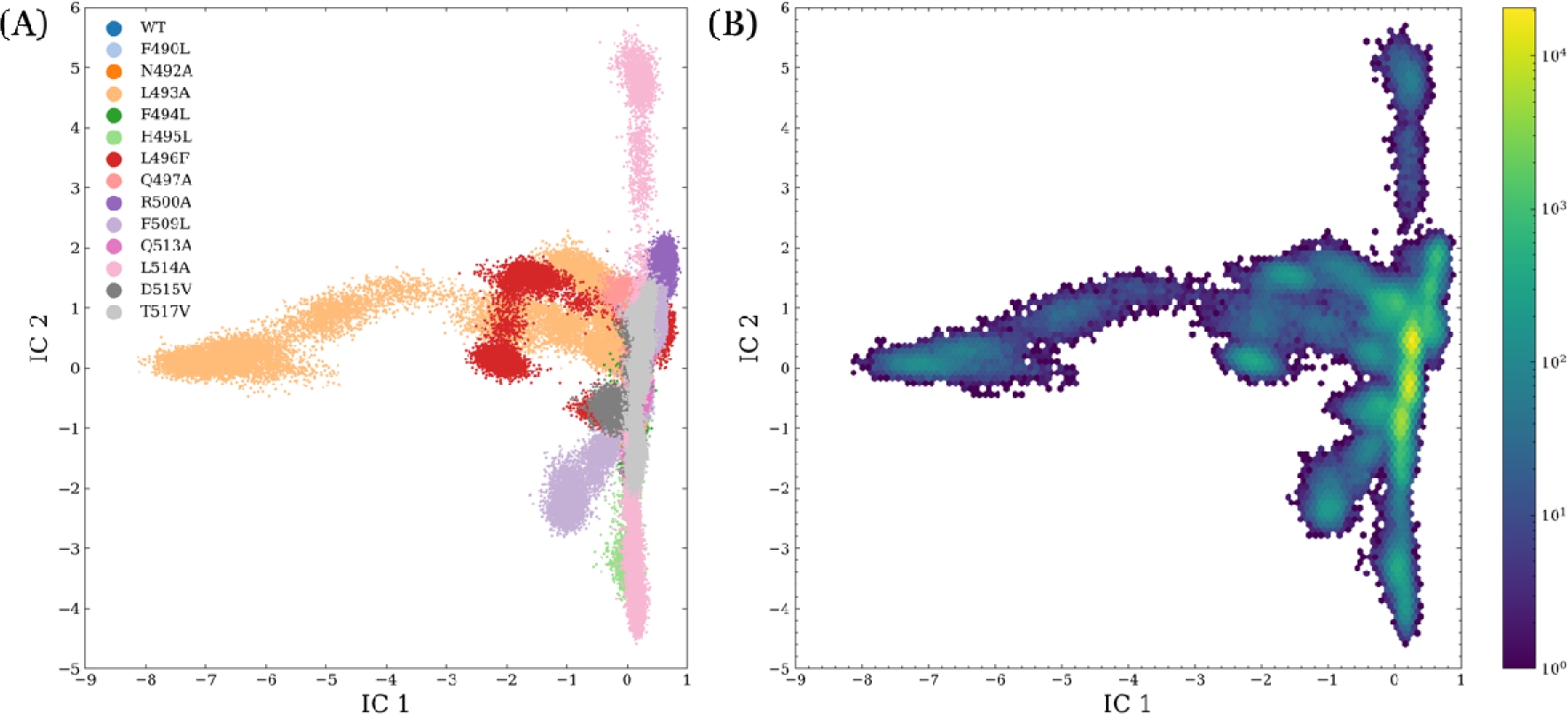
Projection of light states simulation onto two-dimensional space based on t-ICA. **(A)** Distribution of each individual system. **(B)** Overall population of combined light states simulations.

Markov State Model (MSM) is employed to further reveal the dynamic patterns by kinetically clustering the conformations. Through a k-means clustering analysis within the t-ICA 2D space, the simulation data are divided into 50 microstates. The implied timescales were calculated with lag times ranging from 0.1 to 30 ns and the top 15 timescales (Error! Reference source not found.). The trend of the implied relaxation timescale revealed that the estimated timescale converged after about 20 ns, which was chosen as the lag time in the construction of MSM. The differentiation of timescales led us to define seven distinct macrostates for the model. Sequentially, PCCA is used to kinetically cluster these seven macrostates.

All seven macrostates are plotted in the t-ICA 2D space (**Figure 10A**). The population of overall light states simulations in each macrostate is plotted in **Figure 10B**. The macrostates 2 and 1 has the two most dominant occupancies as 78.72% and 11.92%, respectively. The WT light state populates both macrostates 1 and 2, echoing the occupancy patterns illustrated in **Figure 9B**, where these states correspond to the most populated regions. In addition to macrostate 2, certain mutants explored unique conformational space represented as different macrostates (Error! Reference source not found.). Notably, L493A, H495L, L496F, F509L, and L514A mutants distinctly explore macrostates 3 through 7. This diversity in macrostate occupancy provides some details of unique dynamic behaviors of these mutants.

**Figure 10.**
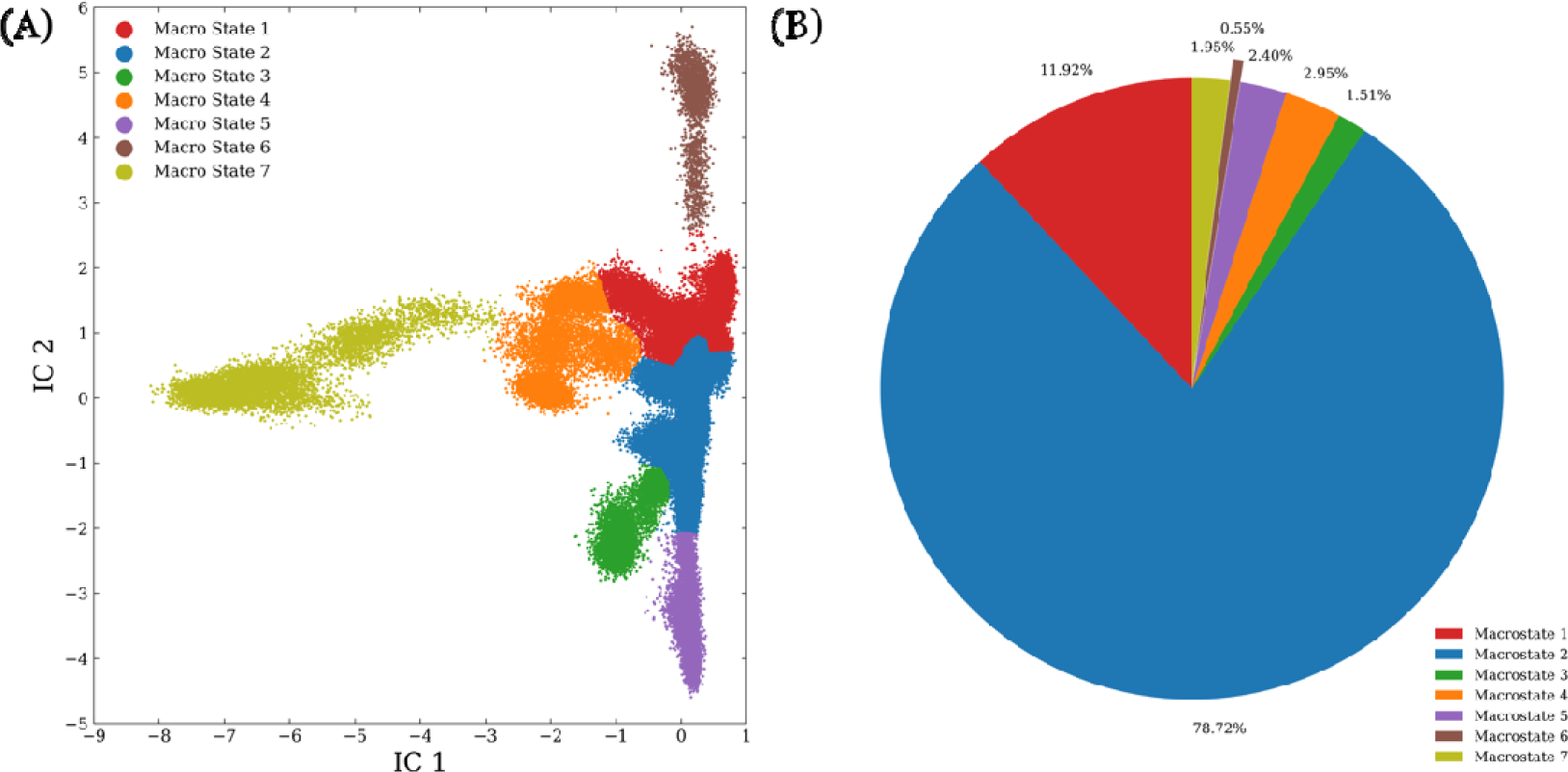
Markov state model of *As*LOV2 light states. **(A)** The macrostates represented on the t-ICA 2D space. **(B)** Relative population composition of the macrostates.

## Discussion

Integrating findings from experimental analyses with computational data, we have chosen four mutants for an in-depth investigation from the pool of thirteen. This selection comprises three mutants located on the Hβ sheet (L493A, L496F, and Q497A) and one on the Iβ sheet (D515V), based on th simulation dynamics, conformation exploration, and experimental results comprehensively. Our focused study aims to uncover the underlying reasons for the distinct behaviors exhibited by these specific mutants. We first extract the correlation values from DCCM for residues surrounding the mutant sites. The criteria to define surrounding residues is that in the first frame of WT or mutant production runs, any residue with at least one atom within 3 Å to any atom in the mutant site (**Figure 11**). Baker-Hubbard method is used to identify hydrogen bonds in the selected region that appear at least 50% of the total frames in three simulations.

**Figure 11.**
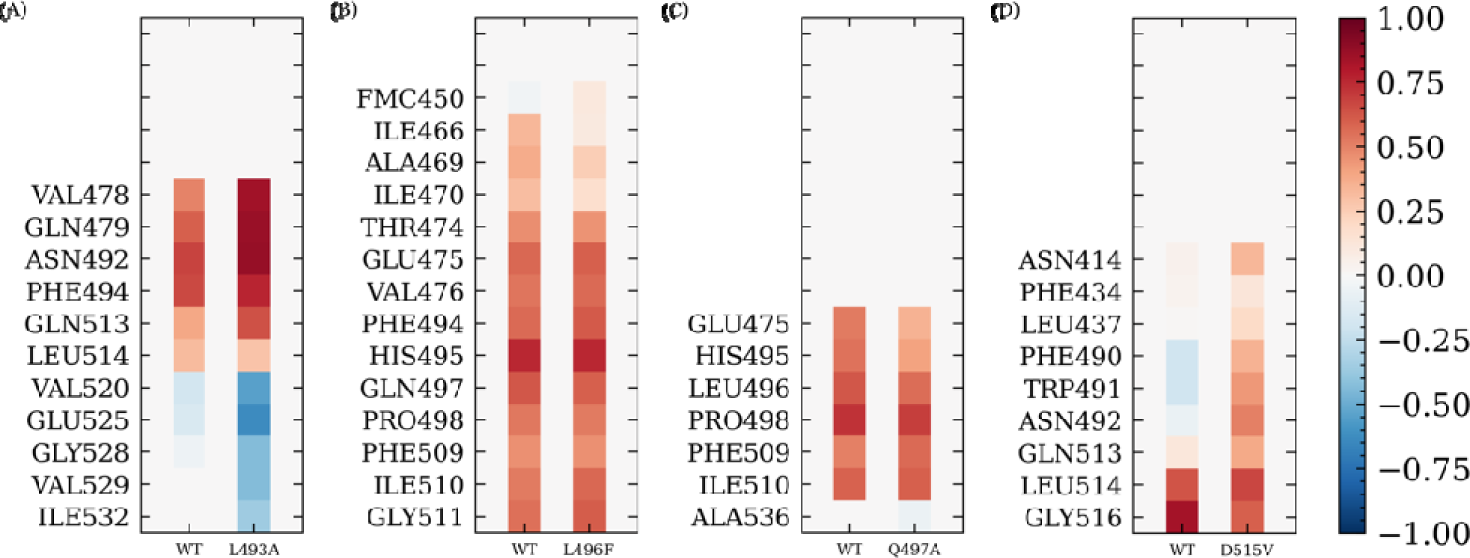
The comparison of correlation between the mutated residue and its adjacent residues in the mutant and in WT for **(A)** L493A, **(B)** L496F, **(C)** Q497A, and **(D)** D515V.

To further investigate and compare the dynamical behavior of these four mutants regarding WT light state, the hydrogen bonds were analyzed (Error! Reference source not found. and **Figure 12**). All four mutants were found to lack the hydrogen bond between Thr407 and Ala452 found in the WT. Thr407 was identified to be essential in the unfolding mechanism of A’α helix^15^. This missing hydrogen bond is suggested to be the main reason of the high fluctuations observed in the N-terminal.

**Figure 12.**
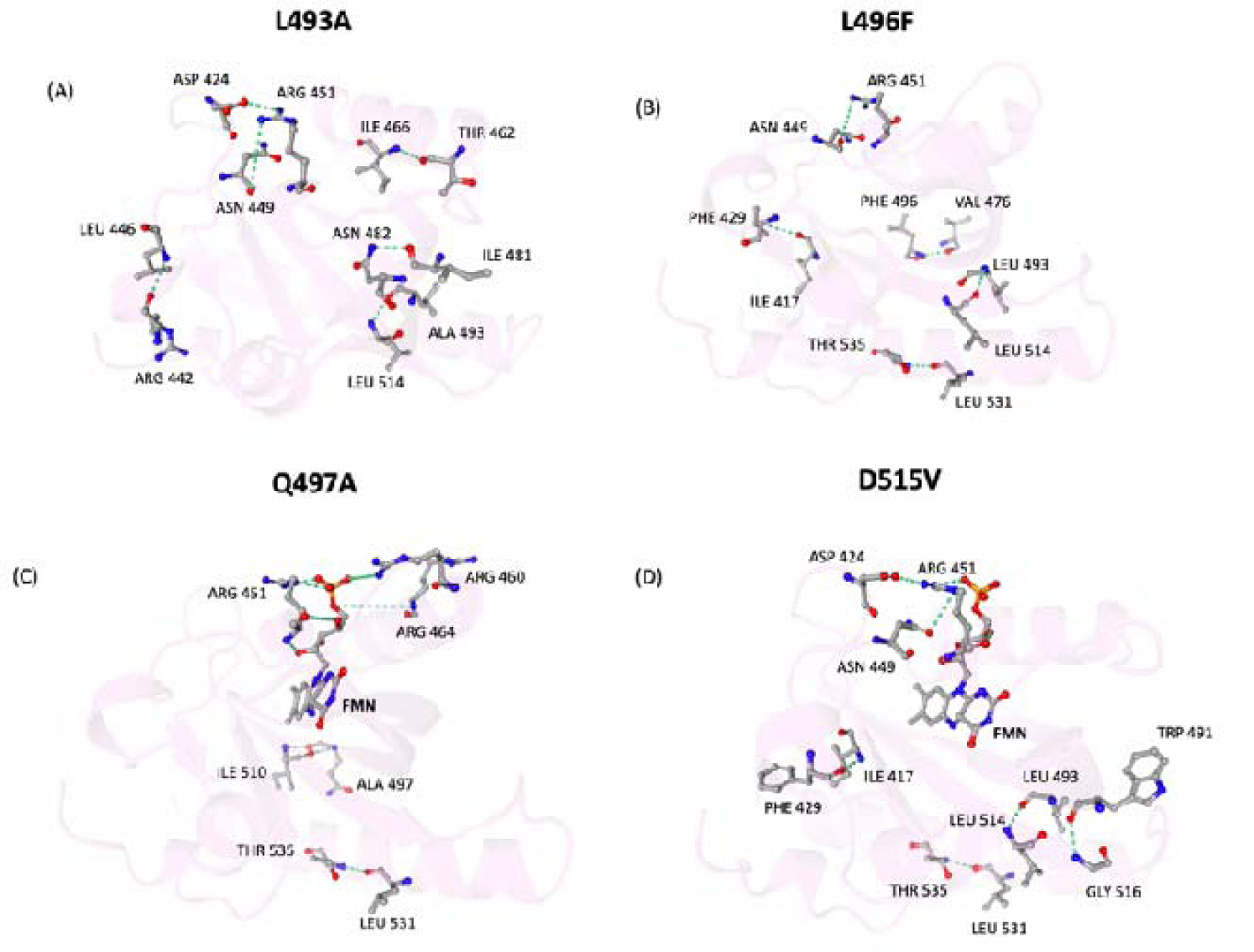
The unique hydrogen bonds patterns formed with the most disruptive mutants compared to WT for **(A)** L493A hydrogen bonds, **(B)** L496F hydrogen bonds, **(C)** Q497A hydrogen bonds, and **(D)** D515V hydrogen bonds.

Compared to WT, we observe several missing hydrogen bonds between flexible secondary structure in L493A mutant. Among these missing hydrogen bonds, two missing two hydrogen bonds between Iβ sheet and A’α helix, one between Ser431-O and GLys413-O and one between Ser433-OG and Glu412-O, could lead to increasing flexibility of the A’α helix. Another two missing hydrogen bonds between Iβ sheet and loop connecting Iβ sheet and Jα helix, one between Gln479-NE2 and Val520-O and one between Gln479-OE1 and Val520-N, may lead to increasing flexibility of the Jα helix. There is another missing hydrogen bond between Gβ sheet and Eα helix as one between Ser486-OG and Glu439-OE2. On the other hand, the L493A mutant has more hydrogen bonds within secondary structures or positive correlated regions than the WT. As shown in the local correlation patterns, residue is having a more negative correlation with Jα helix but a more positive correlation with Gβ sheet (**Figure 11A**). As a result, L493A exhibits a higher flexibility with high fluctuations and the amplified correlations patterns (**Figure 12A**).

For L496F mutant, Gln479-NE2 with Val520-O and Gln479-OE1 with Val520-N are missing compared to the WT, leading to a more flexible Jα helix. The hydrogen bond between Asn482-ND2 on Gβ sheet and Leu453-O on Eα helix are also absent. This leads to a more flexible Eα helix and Fα helix, and it is observed that the correlation between residue 496 to Fα helix are less positive (**Figure 11B**). These absent hydrogen bonds might result from the steric hider of the larger Phenylalanine side chain (**Figure 12B**), which could contribute to the different distribution of L493A and L496F mutants from the WT and other mutants in the t-ICA 2D space (**Figure 9A**).

For Q497A mutant, the two hydrogen bonds between Gln497 and Ile510 remain. In addition, the Cys450-FMN adduct has more hydrogen bonds associated with Arg451, Arg460, and Arg464 on Fα helix (**Figure 12C**). These hydrogen bonds may help the Q497A mutant maintain the local correlation, protein flexibility, and protein dynamics similar to the WT (**Figure 11C**). Similar changes in the hydrogen bonds patterns of the Cys450-FMN core were observed in D515V mutant. This can be attributed that Asn482 form an extra hydrogen bond in both mutants with Leu453 compared to other mutants (**Figure 12D**). This hydrogen bond is disrupting the hydrogen bonds of Asn482 with FMN core in both Q497A and D515V, explaining the drastic changes in the hydrogen bond patterns observed in the FMN core.

For D515V mutant, we observe an increased number of hydrogen bonds. Especially, two hydrogen bonds occur near residue 515: one between Leu493-N and Leu514-O, and one between Gly516-N and Trp491-O. As comparison, the WT has one hydrogen bond between Leu514-N and Ala493-O. In this mutant, residues Leu493 and Leu514, located on two adjacent anti-parallel β-sheets, should have stronger correlation with each other due to two hydrogen bonds. Similarly, the hydrogen bond between Gly516 and Trp491 also helps to maintain or strengthen the positive correlation comparing to the WT . The correlations between the mutated residue 515 and residues 490-492 are more positive than in the WT (**Figure 11D**). As a result, the Iβ sheet, especially the connecting region to Jα helix, is more constrained, which leads to a reduced flexibility and increased higher Iβ sheet ratio. It is possible when aspartic acid changes to leucine at position 515, the negative side chain becomes neutral and the electrostatic attractions from surroundings (opposite to residues 490-492) becomes steric effect and contribute to the attractions of residues 490-492.

## Conclusion

Extensive Molecular dynamics simulations were carried out for the wild type and 13 different mutations introduced to the Hβ and Iβ sheets of *As*LOV2 to investigate the role of β-sheets in the allosteric mechanism of this protein. RMSF, DCCM, and MSM analyses indicated that four mutations were associated with most of the changes. The L493A mutant is associated with the highest fluctuations in different regions of the proteins. The L493A and L496F mutants are among the mutations exploring most of the conformational space likely due to the missing hydrogen bond between Val520 and Gln479 in these two mutants. The Q497A and D515V mutants affect the hydrogen bond patterns formed with the FMN molecule more than any other mutations due to the missing hydrogen bond between Asn482 and Leu453.

## Abbreviations

MD: molecular dynamics
RMSF: root-mean-square fluctuations
DSSP: dictionary of protein secondary structure
DCCM: dynamic cross-correlation matrix
N/A: not applicable.

## Data Availability Statement

The processed results presented or mentioned in this study can be found in Zenodo repository at https://github.com/smu-tao-group/VAE_protein_assessment.

## Author Contributions

**Sian Xiao:** Conceptualization, Methodology, Software, Formal analysis, Investigation, Data Curation, Validation, Visualization, Writing - Original Draft, Writing - Review & Editing. **Mayar Tarek Ibrahim:** Conceptualization, Methodology, Formal analysis, Investigation, Validation, Writing - Original Draft, Writing - Review & Editing. **Gennady Verkhivker:** Writing - Review & Editing. **Brian Zoltowski:** Writing - Review & Editing. **Peng Tao:** Supervision, Project administration, Funding acquisition, Resources, Writing - Review & Editing.

## Acknowledgments

Computational time was provided by the Southern Methodist University’s Center for Research Computing.

