## Supplemental Figures S1-S7, Tables S1,S2 for "Microsecond Molecular Dynamics Simulations and Markov State Models of Mutation-Induced Allosteric Mechanisms for the Light-Oxygen-Voltage 2 Protein : Revealing Structural Basis of Signal Transmission Induced by Photoactivation of the Light Protein State"

### Appendix

#### Theory details of TICA and Markov state model

TICA looks for the slowest-relaxing degree of freedom by solving a generalized eigenvalue problem as:

$$\bar{C}F = CFK$$

where $K = diag(k_1, \cdots, k_n)$ and $F = (f_1, \cdots, f_n)$ are eigenvalue and eigenvector matrices, respectively. $C$ is the covariance matrix, $\bar{C}$ is the time-lagged covariance matrix with a certain lag time $\Delta t$, calculated by:

$$C = \langle (x(t) - \langle x(t) \rangle)(x(t) - \langle x(t) \rangle)^T \rangle$$

$$\bar{C} = \langle (x(t) - \langle x(t) \rangle)(x(t + \Delta t) - \langle x(t) \rangle)^T \rangle$$

where $\langle \cdots \rangle$ denotes the time average.

In order to obtain a symmetric time-lagged covariance matrix, $\frac{1}{2}(\bar{C} + \bar{C}^T)$ is calculated. The latter step assumes the time reversibility of the process, which is satisfied in MD simulations.

The transition matrix and transition probability were calculated to quantify the transition dynamics among macrostates. The corresponding transition probability from state $i$ to state $j$ is calculated as:

$$\mathrm{P_{ij}}(\tau) = \mathrm{Prob}\left(\mathrm{x_t} + \tau \in \mathrm{S_j} \middle| \mathrm{x_t} \in \mathrm{Si}\right)$$

A proper lag time is required for MSM to be Markovian. The value of the lag time and the number of macrostates are selected based on the result of estimated relaxation timescale. The implied time scale can be calculated using the eigenvalues ($\lambda_i$) in the transition matrix as

$$t_i = -\frac{\tau}{\ln|\lambda_i(\tau)|}$$

The number of protein metastable states associated with these slow relaxation timescales can be inferred based on the convergence of implied relaxation time scale. These metastable states effectively discretize the conformational landscape.

#### Average across parallel simulations

The properties are calculated for each independent simulation, including mean RMSF values over certain residues, DSSP ratio, DCCM. Then mean values and standard deviations for these values are calculated. In DCCM plots, mean values are shown. In other bar plots, mean values are represented by the bar heights and the error bars represent the standard deviations. The standard deviation is calculated as the square root of the average of the squared deviations from the mean:

$$std = \sqrt{\overline{(x - \bar{x})^2}}$$

where $x$ represents individual sample and $\bar{x}$ represent the averaged values.

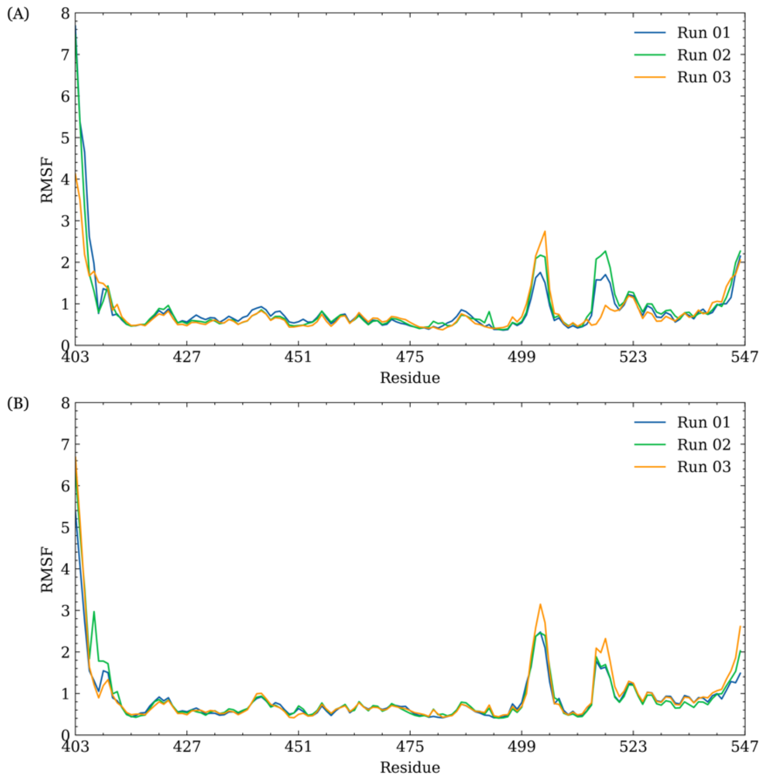


**Figure S1.** Root-mean-square fluctuations (RMSF) for WT *As*LOV2 in **(A)** dark state and **(B)** light state in three parallel simulations.

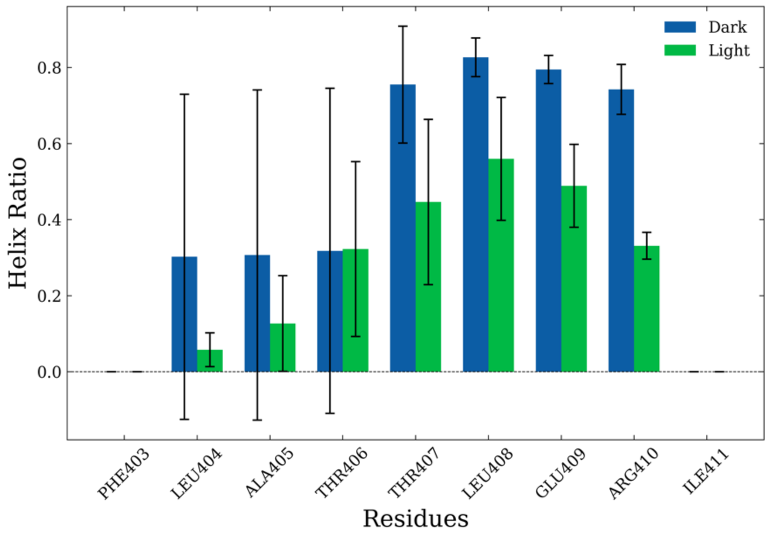


**Figure S2.** Ratio of helical content of the residues on N-terminal. The average values and error bars are calculated based on three parallel simulations.

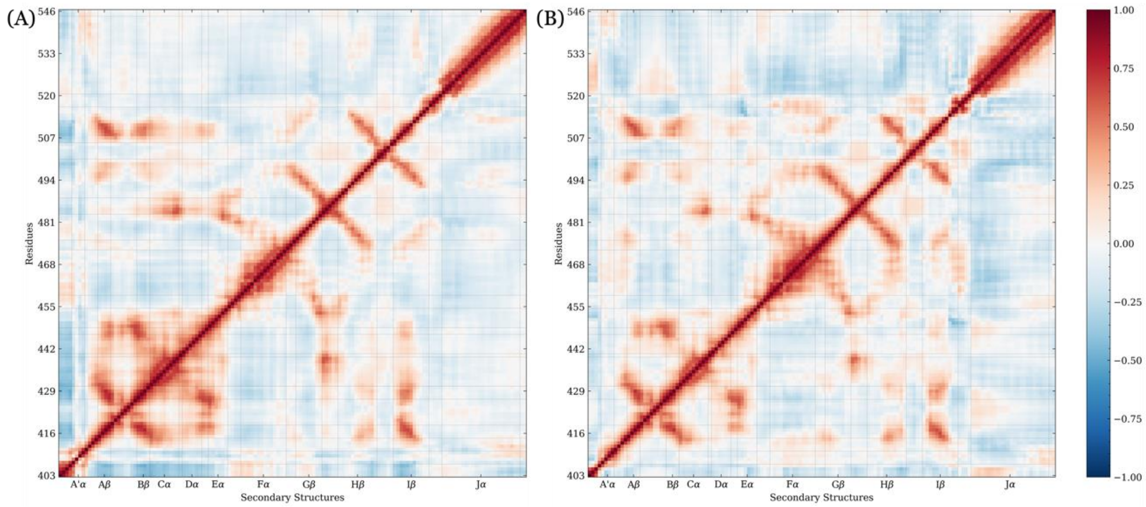

**Figure S3.** Average dynamic cross-correlation matrix for *As*LOV2 WT in **(A)** dark state and **(B)** light state over three parallel simulations.

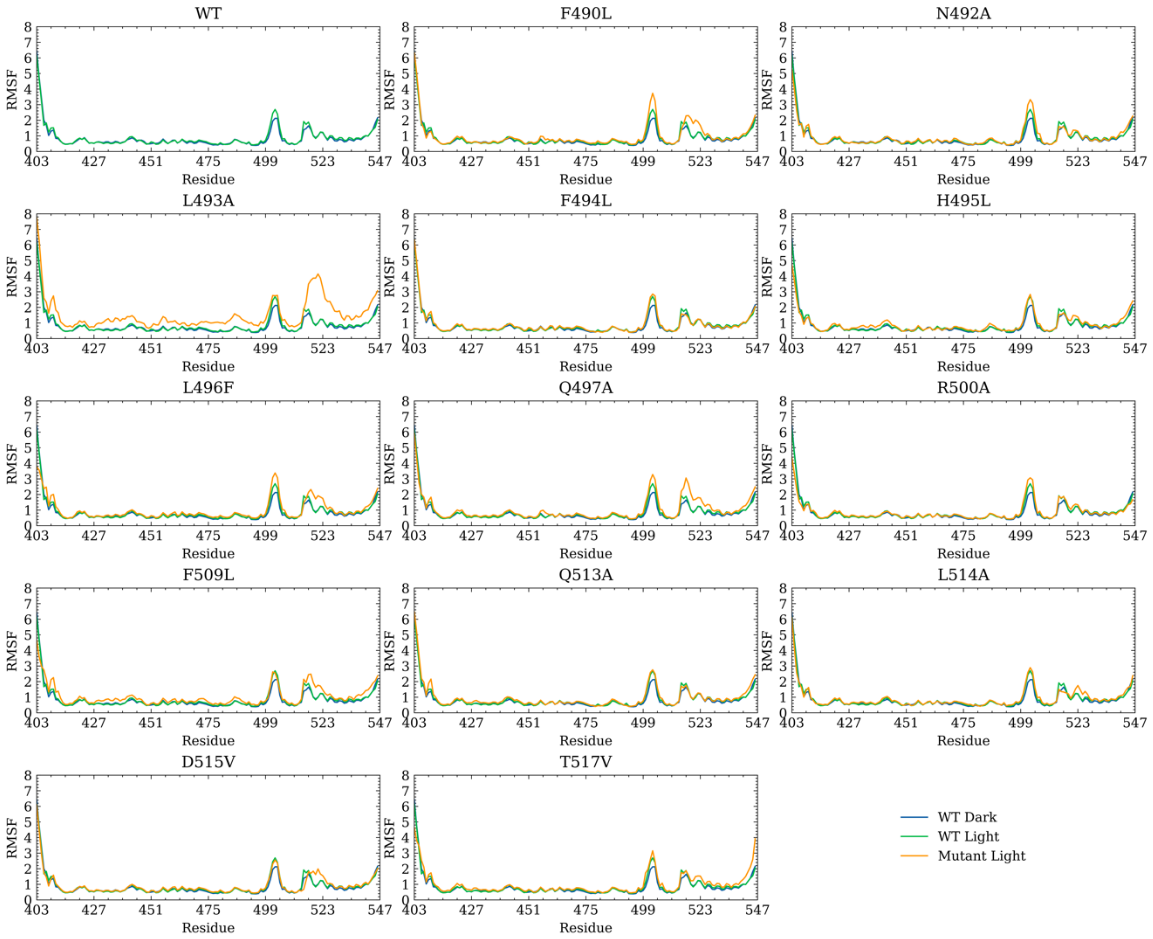

**Figure S4.** Average root-mean-square fluctuations (RMSF) of WT (dark state in blue and light state in green) and mutants (light state in orange) over three parallel simulations.

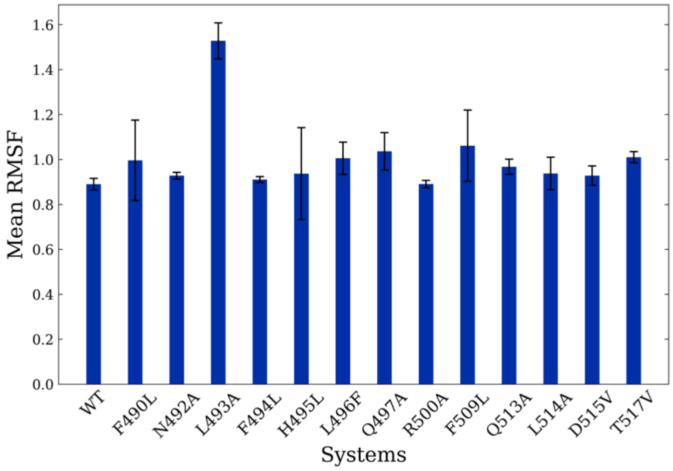


**Figure S5.** Average RMSF values over the whole protein of WT and mutants in light state.

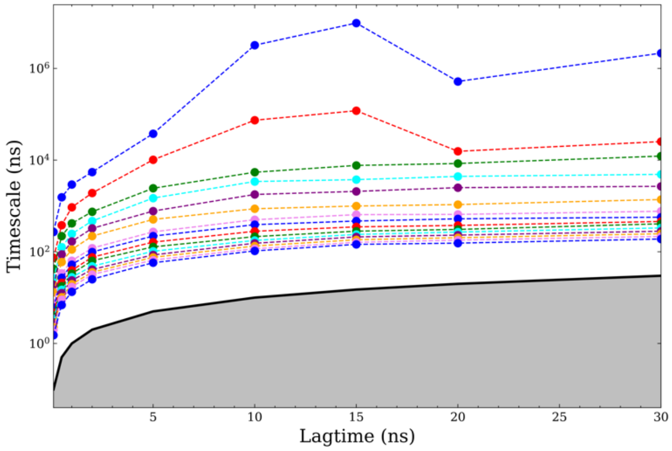


**Figure S6.** The estimated relaxation timescale is based on the transition probabilities among different microstates using different lag times ranging from 0.1 to 30 ns. The top 15 timescales are presented. The black line indicates where the lag time equals the relaxation timescale. Implied timescales intersecting this gray area denote processes faster than the lag time, suggesting that these processes are too rapid to be resolved.

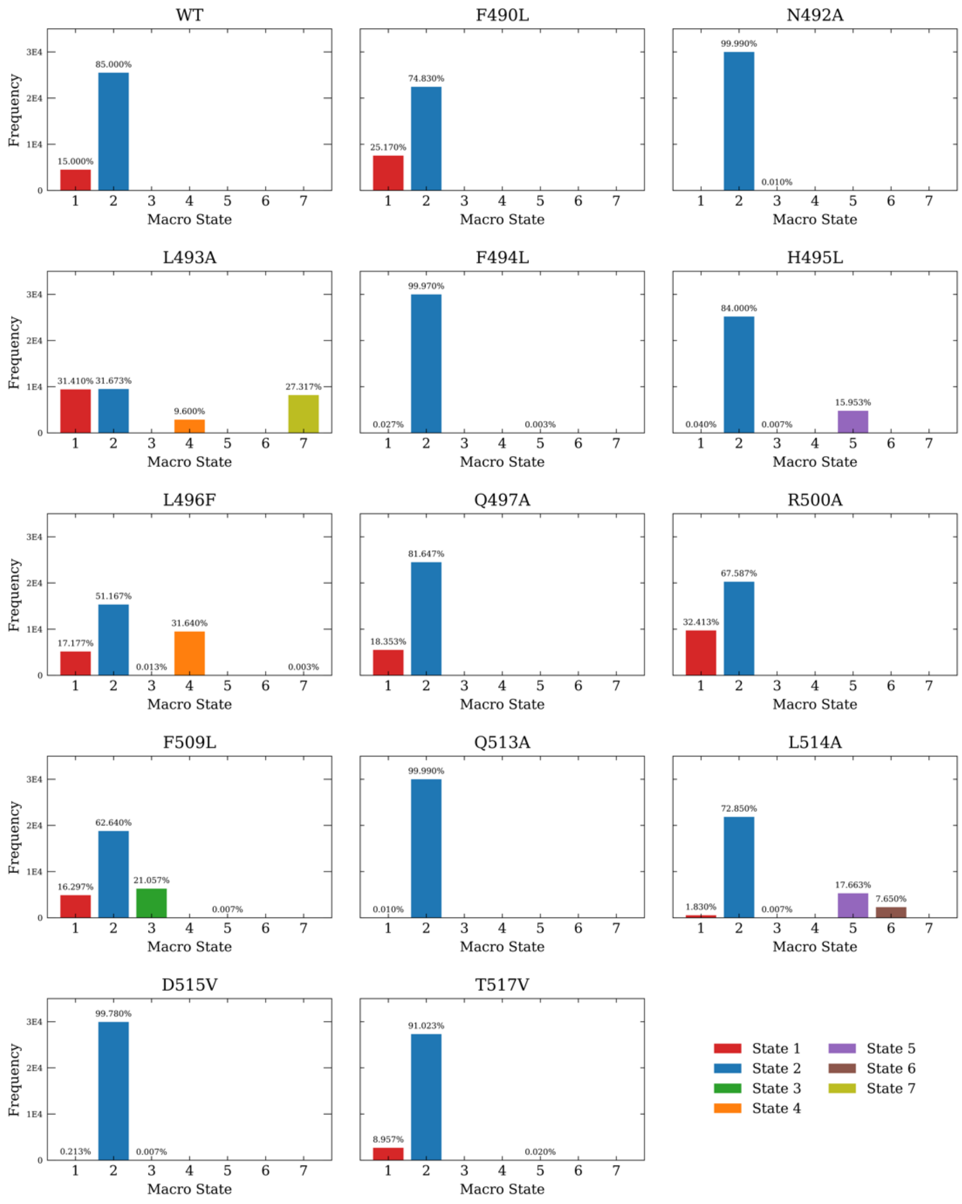


**Figure S7.** Macrostate occupancy profiles for different systems including WT and mutants.

### Tables

**Table S1.** Most common secondary structure configuration assigned by Dictionary of protein secondary structure (DSSP) during three parallel simulations for the N-terminal. The A'α helix residues are noted in italic.

| **State** | **PHE403** | **LEU404** | **ALA405** | ***THR406*** | ***THR407*** | ***LEU408*** | ***GLU409*** | **ARG410** | **ILE411** |
|---|---|---|---|---|---|---|---|---|---|
| Dark | CCC | CCH | CCH | *CCH* | *HHH* | *HHH* | *HHH* | HHH | CCC |
| Light | CCC | CCC | CCC | *CCH* | *HCH* | *HCH* | *HCH* | CCC | CCC |

**Table S2.** Hydrogen bonds occurrence comparison between WT and mutants. The hydrogen bond (donor & acceptor) is present in this state but missing in the other.

| **L493A** | **WT** | **L496F** | **WT** |
|---|---|---|---|
| PHE429-N &ILE417-O | THR407-OG1&ALA542-O | PHE429-N &ILE417-O | THR407-OG1&ALA542-O |
| LEU446-N &ARG442-O | ARG421-N &ASP419-OD1 | ARG451-N &ASN449-OD1 | ARG421-N &ASP419-OD1 |
| ARG451-N &ASN449-OD1 | ARG421-NE &ASP419-OD1 | ARG451-NH1&ASP424-OD2 | ARG451-NH1&ASP424-OD1 |
| ARG451-NH1&ASP424-OD2 | SER431-OG &LYS413-O | VAL476-N &PHE496-O | ASN472-ND2&ASP468-O |
| ILE466-N &THR462-O | SER433-OG &GLU412-O | LEU493-N &LEU514-O | VAL476-N &LEU496-O |
| ASN482-ND2&ILE481-O | GLN454-NE2&FMC450-O4' | PHE496-N &VAL476-O | GLN479-NE2&VAL520-O |
| LEU514-N &ALA493-O | GLN479-NE2&VAL520-O | THR535-OG1&LEU531-O | ASN482-ND2&LEU453-O |
| THR535-OG1&LEU531-O | ASN482-ND2&LEU453-O | | LEU496-N &VAL476-O |
| | SER486-OG &GLU439-OE2 | | VAL520-N &GLN479-OE1 |
| | LEU514-N &LEU493-O | | |
| | VAL520-N &GLN479-OE1 | | |
| | MET530-N &ARG526-O | | |

| **Q497A** | **WT** | **D515V** | **WT** |
|---|---|---|---|
| FMC450-O4'&FMC450-O3P | THR407-OG1&ALA542-O | PHE429-N &ILE417-O | THR407-OG1&ALA542-O |
| ARG451-NE &FMC450-O3P | ARG421-N &ASP419-OD1 | FMC450-O4'&FMC450-O3P | ARG421-N &ASP419-OD1 |
| ARG451-NH2&FMC450-O2P | ARG421-NE &ASP419-OD1 | ARG451-N &ASN449-OD1 | ARG421-NH2&ASP419-OD1 |
| ARG460-NH1&FMC450-O3P | ARG421-NH2&ASP419-OD1 | ARG451-NE &FMC450-O3P | ARG421-NH2&ASP419-OD2 |
| ARG460-NH2&FMC450-O1P | ARG421-NH2&ASP419-OD2 | ARG451-NH1&ASP424-OD2 | ASN472-ND2&ASP468-O |
| ARG464-NH1&FMC450-O1P | ASN492-ND2&ASN482-OD1 | ARG451-NH2&FMC450-O2P | |
| ALA497-N &ILE510-O | GLN497-N &ILE510-O | LEU493-N &LEU514-O | |
| ILE510-N &ALA497-O | ASP501-N &ASP505-O | GLY516-N &TRP491-O | |
| THR535-OG1&LEU531-O | GLY504-N &ASP501-O | THR535-OG1&LEU531-O | |
| | ILE510-N &GLN497-O | | |
